# Maximum Parsimony Inference of Phylogenetic Networks in the Presence of Polyploid Complexes

**DOI:** 10.1101/2020.09.28.317651

**Authors:** Zhi Yan, Zhen Cao, Yushu Liu, Luay Nakhleh

## Abstract

Phylogenetic networks provide a powerful framework for modeling and analyzing reticulate evolutionary histories. While polyploidy has been shown to be prevalent not only in plants but also in other groups of eukaryotic species, most work done thus far on phylogenetic network inference assumes diploid hybridization. These inference methods have been applied, with varying degrees of success, to data sets with polyploid species, even though polyploidy violates the mathematical assumptions underlying these methods. Statistical methods were developed recently for handling specific types of polyploids and so were parsimony methods that could handle polyploidy more generally yet while excluding processes such as incomplete lineage sorting. In this paper, we introduce a new method for inferring most parsimonious phylogenetic networks on data that include polyploid species. Taking gene trees as input, the method seeks a phylogenetic network that minimizes deep coalescences while accounting for polyploidy. The method could also infer trees, thus potentially distinguishing between auto- and allo-polyploidy. We demonstrate the performance of the method on both simulated and biological data. The inference method as well as a method for evaluating given phylogenetic networks are implemented and publicly available in the PhyloNet software package.

## 1 Introduction

Hybridization and polyploidization have long been recognized as crucial factors in speciation and genomic and phenotypic novelties [24, 2]. While in homoploid hybridization, the hybrid has the same number of chromosome sets as the two parental species, allopolyploid hybrids receive both chromosome sets from the parents, thus increasing the size of the chromosome set as compared to the two parents. Both types of hybridization result in reticulate evolutionary histories that are best modeled by phylogenetic networks. Autopolyploidy, on the other hand, is whole-genome duplication (WGD) that involves a single lineage and does not violate a treelike evolutionary history.

Polyploidy is prevalent across the eukaryotic branch of the Tree of Life. Most of the extant flowering plants are polyploids, and many of the other diploid plants can be traced back to one or more rounds of ancient WGD events [19, 10]. Although hybrid and polyploid species are less commonly observed in animals than plants, presumably owing to their potential reduced fitness, fish and amphibians are known to have high incidence of polyploidy [7, 1, 33]. Furthermore, it is believed that at least two rounds of ancient WGD occurred in the vertebrate lineage (the 2R hypothesis) [23, 20]. Fungi also include polyploids, with evidence supporting that the ancestor of the baker’s yeast *Saccharomyces cerevisiae* underwent WGD [18].

From modeling and inference perspectives, allopolyploidy is of particular interest, as it results from hybridization of two species and gives rise to evolutionary histories in the form of phylogenetic networks. Although polyploidy could potentially be identified from the chromosome count, the task of determining its mode of origin is non-trivial, especially when the parental taxa are closely related. Moreover, other evolutionary processes, such as incomplete linaege sorting (ILS), complicate this task. Therefore, extending models such as the multispecies coalescent to account for polyploidy could provide a powerful approach inferring evolutionary histories of polyploids.

[36, 37] extended the multispecies coalescent to incorporate hybridization into the model, giving rise to the multispecies network coalescent (MSNC). Based on this generative process, PhyloNet [28, 32] implements a wide array of methods for inferring phylogenetic networks in the presence of incomplete lineage sorting and diploid hybrids, including parsimony methods [35], maximum likelihood and pseudo-likelihood methods [37, 38, 41], and Bayesian methods [30, 31, 42]. Since almost all these methods are computationally demanding (with the exception of [38]), a divide-and-conquer approach was recently introduced to speed them up [39]. [3] illustrated the use of many of these inference methods, as well as network summarization methods, on data generated under the multispecies network coalescence, which assumes diploid hybridization.

[12] evaluated various inference tools in PhyloNet [28, 32] for inferring phylogenetic networks of polyploid strawberries, *Fragaria* (Rosaceae), with species that ranged in ploidy from tetraploid to decaploid. As [2] correctly pointed out, “Though not specifically built for polyploids, PhyloNet can model multiple haplotypes in each lineage. It can hence infer hybridization in these species when the assumptions of its model (the coalescent) are not violated, making it most appropriate for recent allopolyploids.”

While existing inference tools in PhyloNet were not designed for handling polyploidy, there are several existing methods that are designed specifically to model polyploidy events [24]. Many of these methods have relied on multi-labeled trees, or MUL-trees, to model polyploids. As multiple copies of a locus could be present in the genome due to polyploidization, a MUL-tree extends standard phylogenetic trees by allowing multiple leaves to be labeled by the same taxon name (figure 1a-d). There is a straightforward connection between a phylogenetic network and a MUL-tree, as illustrated in figure 1a-b. Indeed, in one of the earliest works in this area, [8] provided an algorithm for converting a MUL-tree into a phylogenetic network, which was later implemented in the PADRE software [15]. This connection between phylogenetic networks and MUL-trees was the basis for computations under the MSNC in [36, 37] before moving towards computations directly on the phylogenetic network in subsequent implements in PhyloNet. The software tool GRAMPA [29] uses MUL-trees to reconcile a set of gene trees parsimoniously with a given species tree to postulate polyploidy events.

**Figure 1:**
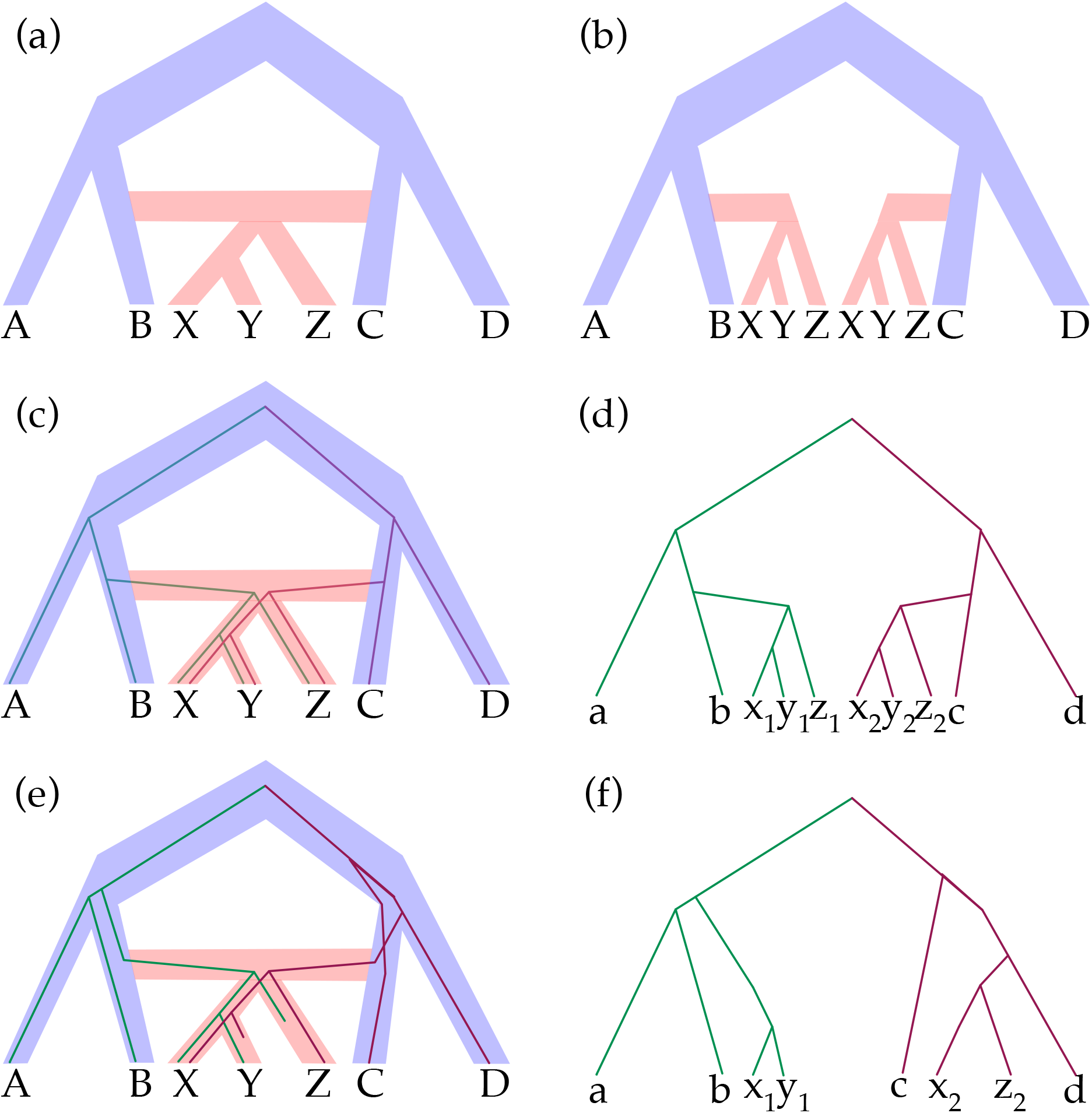
Allopolyploidy, phylogenetic networks, and MUL-trees. (a) Phylogenetic network depicting allopolyploidization involving the ancestor of *X, Y*, and *Z*. (b) MUL-tree representation of the network. (c) Gene tree inside the branches of the phylogenetic network. (d) Gene tree with two copies of the gene in each of the species *X, Y*, and *Z*. (e) Gene tree inside the branches of the phylogenetic network in the presence of incomplete lineage sorting and gene loss. (f) Gene tree where the signal for the allopolyploids is confounded by ILS and gene loss.

The only statistical inference methods specifically designed for handling allopolyploids that we are aware of are the AlloppNET method of [11] and its extension [24]. AlloppNET uses Bayesian Markov chain Monte Carlo (MCMC) to sample, using multi-locus DNA sequence data, the posterior distribution of phylogenetic networks that contain diploid and allotetraploid species. The method allows for multiple individuals per species and samples, in addition to the phylogenetic network topology, parameters including divergence and hybridization times as well as population sizes. Like other statistical inference methods in PhyloNet, AlloppNET is computationally intensive, in particular as it employs reversible-jump MCMC to sample the trans-dimensional space of phylogenetic networks. Furthermore, as described in [2], producing an all-encompassing stochastic model of polyploidization would be a massive undertaking due to the complexities of processes occurring during and after polyploidization events.

Recognizing that the parsimony method of [35] as implemented in PhyloNet was not designed to handle allopolyploids, [22] coupled it with a permutation scheme where multiple analyses are conducted in each of which only two copies from the polyploid are mapped to one parent and the other copies are mapped to a second parent, and then reporting the optimal result over all these analyses. In this work, we extend the method of [35] to properly handle polyploids without the need for a permutation approach. We implemented both an inference method and a scoring method in PhyloNet and assessed their performance on the simulated and biological data used in [22]. We report on the accuracy and running time of the new method, and compare it to existing methods in PhyloNet that were not designed to handle polyploid hybridization.

## 2 Methods

### 2.1 Minimizing Deep Coalescences in the Presence of Allopolyploidy

In his seminal paper, [16] proposed minimizing deep coalescences (MDC) as a parsimony criterion for reconciling a gene tree with a given species tree, as well as for inferring a species tree from a collection of gene trees, both under the assumption that gene tree heterogeneity is caused by ILS. [17] later implemented and tested a heuristic for inferring a species tree under the MDC criterion. [27] provided a mathematical characterization of the number of deep coalescences given a clade in the species tree (without having the species tree itself), which allowed for developing exact algorithms for species tree inference under the MDC criterion. To account for hybridization and introgression simultaneously with ILS, [35] introduced the MDC criterion for inferring phylogenetic networks from a collection of gene trees whose heterogeneity is assumed to be caused by ILS and introgression. All these works on the MDC criterion naturally allow for including multiple individuals per species.

While the method of [35] could in practice be used on data with allopolyploid species while treating multiple gene copies as alleles from different individuals, the criterion is not mathematically designed for handling polyploids. Let us illustrate this issue with the scenario depicted in figure 1e. The gene tree shown inside the branches of the phylogenetic network has two copies from each of the three taxa X, Y, and Z, due to the polyploid hybridization event involving taxa B and C. If each of the two copies are treated as alleles from two different individuals, then this gene history will be heavily penalized by the MDC criterion as lineages failed to coalesce on the four branches connected to X, to Y, to Z, and to the most recent common ancestor (MRCA) of X and Y. Indeed, the MDC score of this gene tree given the phylogenetic network is 4 in this case. However, when considering that this is a polyploid hybridization event and the two lineages correspond to two different copies of the gene, these lineages should not be expected to coalesce on these four branches, and the true score of this phylogenetic network / gene tree reconciliation should be 0; that is, this is a gene tree that “perfectly” fits the species evolutionary history with no deep coalescence events. It is this issue that led [22] to correctly not use the method of [35], rather applying a permutation scheme to it so as to sample gene copies and not leave them in the analysis and treat them as alleles from different individuals of the species. We now describe a modification to the MDC criterion so that it properly handles polyploid hybridization events. This extension can be viewed as a generalization of [29] by allowing for ILS.

Using the notation of [9], we denote by *U*(Ψ) a MUL-tree representation of phylogenetic network Ψ, and we denote by *F*(*T*) a phylogenetic network that corresponds to MUL-tree *T*. As [9] showed, neither *U*(Ψ) nor *F*(*T*) are unique in general, though they are unique for special classes of phylogenetic networks. While the leaves of a gene tree are uniquely mapped to the leaves of a phylogenetic network (since each species labels exactly one leaf in the network), this is not the case for MUL-trees. For example, for the gene tree in figure 1d and the MUL-tree of figure 1b, each of the two copies *x*_1_ and *x*_2_ could map to either of the two leaves labeled by X in the MUL-tree. Figure 2a shows all possible allele mappings for these gene trees and MUL-tree. Given the set 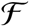 of all allele mappings of a gene tree *g* to a MUL-tree *T*, the number of extra lineages of *g* given *T* is

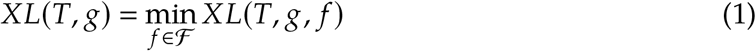

**Figure 2:**
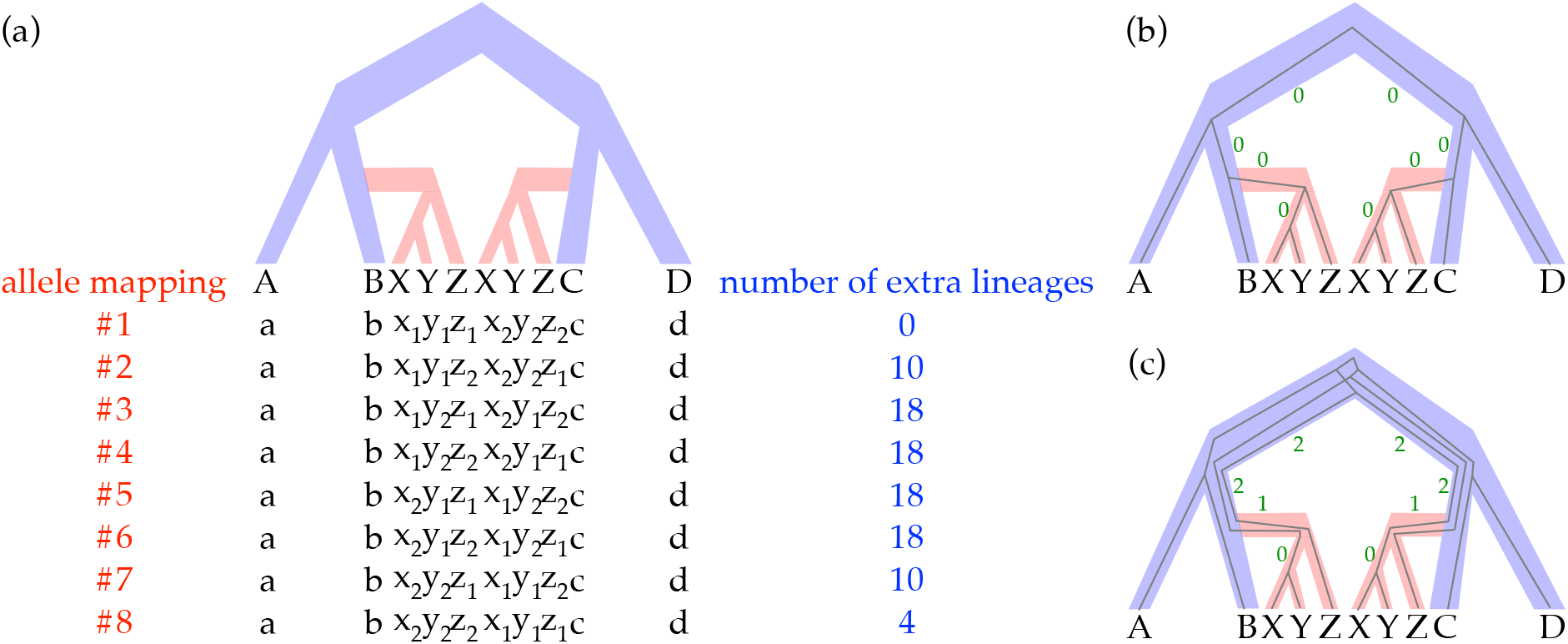
Allele mappings from a gene tree to a MUL-tree and the number of extra lineages. (a) The set of all eight possible allele mappings for the gene tree of figure 1d and the MUL-tree of figure 1b, along with the number of extra lineages that results from each of the allele mappings. (b-c) The reconciliations that correspond to allele mappings #1 and #7, respectively, of the gene tree within the branches of the MUL-tree. Shown next to each of the MUL-tree branches is the number of extra lineages on that branch, which is the number of lineages that persist without coalescing on that branch minus 1.

Here, *XL*(*T, g, f*) is the number of extra lineages given a specific allele mapping *f*, which is the sum, over all branches of the MUL-tree, of the number of extra lineages on each branch given the allele mapping *f*. The number of extra lineages resulting from each of the allele mappings in figure 2a is shown in the same panel, and illustrations of how these quantities are computed for two of the allele mappings are shown in figure 2b-c.

Given a collection 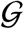 of gene trees, inferring a phylogenetic network Ψ* under the criterion of minimizing the number of extra lineages amounts to computing

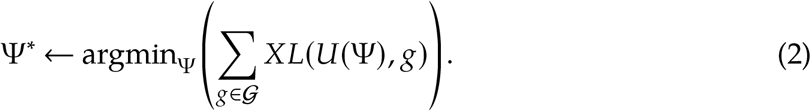

Based on this formulation, the inference is done by walking the space of phylogenetic networks using the implementation of [35], while evaluating the number of extra lineages on the MUL-tree representation of each network, using equation (1).

### 2.2 Handling Autopolyploidy

Unlike allopolyploidy, autopolyploidy is whole-genome duplication that does not involve hybridization between two different species. Therefore, the evolutionary history of a set of species including autopolyploids is a species tree, rather than a network. In this case, a clade whose MRCA in the species tree involves the autopolyploidization event is duplicated in the MUL-tree, yet the two copies are not attached to different locations in the tree, but rather as two sibling clades. The embedding of a gene tree within the branches of the species tree and its corresponding MUL-tree is done as before. These concepts are illustrated in figure 3.

**Figure 3:**
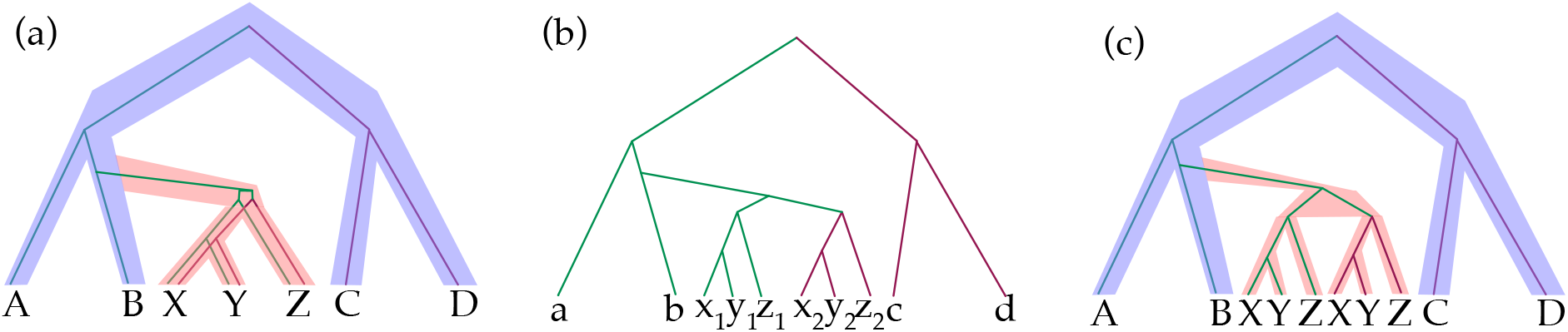
Autopolyploidy and species trees. Autopolyploidy does not violate the treelike structure of the species phylogeny, as it does not involve hybridization between two different species. (a) Species tree with a whole genome duplication involving the ancestor of *X, Y*, and *Z*. (b) Gene tree with both copies of the gene from *X, Y*, and *X*. (c) MUL-tree representation of the species tree, along with the gene tree embedding within its branches.

In this case, inference based on equation (2) can be done in one of two ways. If the event is known to be autopolyploidization, then the search in the network space can be restricted to networks with 0 reticulations—i.e., species trees. If the question being investigated is whether the event is auto- or allopolyploidization, then the search can be conducted while allowing for reticulations. While a phylogenetic network Ψ formed by adding reticulations to an underlying species tree *S* can never have a worse score according to equation (1), the search can still return a species tree, since adding reticulations does not improve the score, or it returns a network whose score can be contrasted to the score of the underlying species tree. In the former case, the method indicates that the event is an autopolyploidization. In the latter case, the difference in scores of the network and underlying tree can be evaluated (albeit not in a principled, statistical manner) to determine the type of polyploidization.

### 2.3 PhyloNet Implementation

We implemented the inference based on equation (2) in PhyloNet [28, 32] as a new command called `InferNetwork_MP_Allopp`. This method takes as input a set of gene trees, and the output is a phylogenetic network (a special case of which is a species tree). In terms of the hybrids in the data, the user can specify two aspects:

- The maximum number of reticulations allowed during the search. If this number is set at 0, then the method searches the space of species trees only. In this case, only autopolyploidization is accounted for.
- Whether the hybrid species are known. If the hybrid species are known and specified by the user, and the maximum number of reticulations allowed equals the number of specified hybrid species, the method searches only networks that have the specified hybrids; that is, it does not detect any other potential hybrids. If the maximum number of reticulations allowed is greater than the number of specified hybrid species, the method could detect additional hybrids. If the hybrids are not specified, then the method identifies hybrids.

In addition to phylogenetic network inference, we provide the user with the option to evaluate, rather than infer, competing hypotheses. Given a phylogenetic network Ψ and a set of gene trees 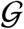, the parsimony score of the Ψ is given by

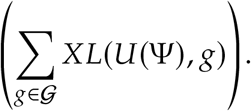

This analysis is enabled by the new PhyloNet command `DeepCoalCount_AlloppNet`. We envision this command used in at least two contexts. First, if the user has two or more evolutionary hypotheses (in the form of phylogenetic networks), this command can be used to assess which of these hypotheses has the best score. Second, if the inference method above returns a phylogenetic network that does not match some biological knowledge, the user can manipulate the inferred network to obtain one that matches the biological knowledge and compare the two. In other words, this command can be used in an exploratory mode.

Last but not least, a brief explanation of the nature of heuristic searches is in order. Inferring the optimal phylogenetic network according to equation (2) is computationally very demanding. Therefore, the algorithm implemented by the command `InferNetwork_MP_Allopp` performs a random walk in the space of phylogenetic networks, evaluates phylogenetic networks encountered during the walk, and returns the best (i.e., the one with the lowest parsimony score) network among all those encountered. This type of heuristics is not guaranteed to find the optimal network, which is why it is recommended to run the command multiple times and return the optimal solution among all the runs (in the simulation experiments below, we ran the method 20 times on each data set).

### 2.4 Simulations

We used the simulated data of [22], which were generated on three evolutionary scenarios, shown in figure 4. The data were generated under three different evolutionary scenarios, where for each scenario population sizes and divergence times were varied to take on three and four different values, respectively, for a total of 12 conditions per scenario. Furthermore, each scenario involved an allopolyploidization event that gave rise to a tetraploid taxon. In total, there were 12 model conditions per scenario, in which one reticulation event occurs in three different scenarios. For each model condition, the original data has 1000 gene trees, and we divided them into 10 replicate data sets, each of which contains 100 gene trees. To study the impact of the number of loci on the relative performance, we used 10, 50 and 100 gene trees of each model condition to infer the species network. Thus, in total we had 3 (scenarios) × 12 (settings per scenario) × 3 (numbers of loci) × 10 (replicates) = 1080 data sets that each method was run on.

**Figure 4:**
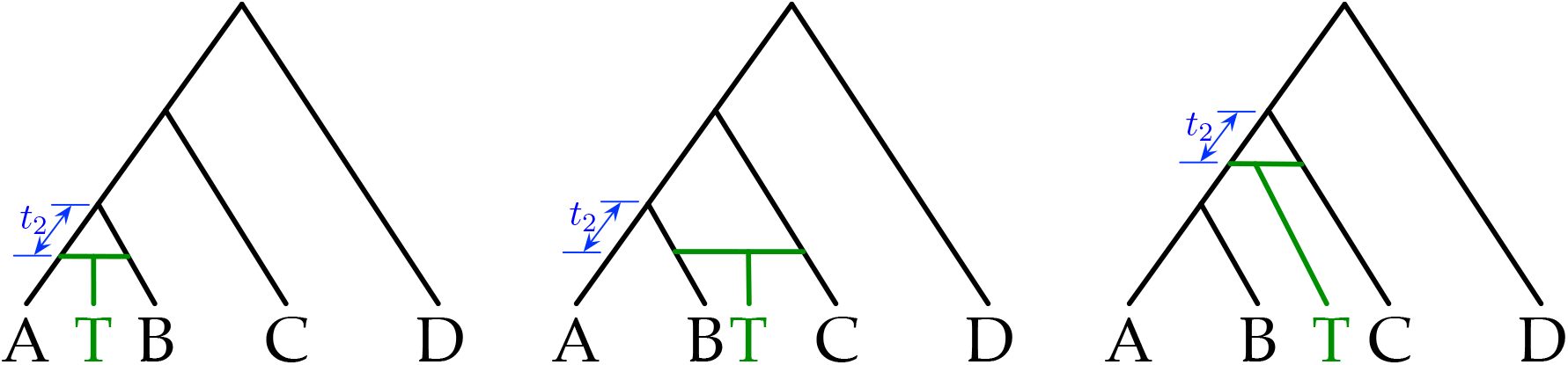
The three evolutionary histories used for simulations in [22]. The scenarios differ in terms of the parents of the tetraploid T. (left) Scenario 1 involves hybridization between two extant, sister taxa. (middle) Scenario 2 involves hybridization between two extant, non-sister taxa. (right) Scenario 3 involves hybridization between two non-sister taxa, one of which is ancestral. Time interval *t*_2_ captures the time between the hybridization event and the most recent divergence event that preceded it.

#### 2.4.1 Evaluating Inferences

We evaluated the accuracy of the methods in terms of the number of correct inferences as well as in terms of the error rates of the inferences [40, 4], which we review here for the sake of completeness.

Let Ψ_*t*_ and Ψ_*i*_ be the true and inferred phylogenetic network topologies, respectively. The backbone distance *BD*(Ψ_*t*_, Ψ_*i*_) is defined as

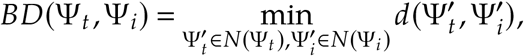

where *N*(Ψ_*t*_) and *N*(Ψ_*i*_) are the sets of all backbone networks of Ψ_*t*_ and Ψ_*i*_, respectively, and *d*(.,.) is the metric of [21]. Finally, letting *t* denote the number of tree nodes (nodes that have a single parent) in the true network, and *r* (Ψ) denote the number of reticulation nodes (nodes that have two parents) in network Ψ, we define:

- True positives: 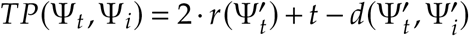.
- True positives rate: 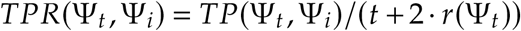.
- False positives: 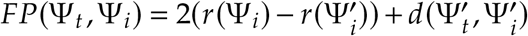.
- False positives rate: *FPR*(Ψ_*t*_, Ψ_*i*_) = *FP*(Ψ_*t*_,Ψ_*i*_)/(*t* + 2 · *r*(Ψ_*i*_)).
- True negatives rate: *TNR*(Ψ_*t*_, Ψ_*i*_) = 1 – *FPR*(Ψ_*t*_,Ψ_*i*_).

### 3 Results and Discussion

#### 3.1 Results on Simulated Data

We tested the new method as well as existing methods in PhyloNet on the 5-taxon simulated data described above. In all the results shown and discussed below, the hybrid species was specified to the methods. Furthermore, as we discussed above, due to the non-deterministic nature of the search heuristics underlying all methods used here, each method was run 20 times on each data set and the optimal solution across all 20 runs was returned.

##### 3.1.1 Performance of MSNC-based Methods: Treating Gene Copies as Alleles from Different Individuals

We ran two existing methods in PhyloNet for inferring phylogenetic networks under the MSNC model: `InferNetwork_ML`, which implements the maximum likelihood method of [37], and `InferNetwork_MPL`, which implements the maximum pseudo-likelihood method of [38] (this is labeled MPL below). For `InferNetwork_ML`, we ran the method in two different modes: On the gene tree topologies as input (this is labeled ML below), and on the gene trees with branch lengths as input (this is labeled as ML_bl below).

As we discussed above, neither of these two methods were designed to handle polyploid species, and they both assume that a gene is present in at most one copy in each individual. Similar to how analyses were conducted using PhyloNet methods in [12], we treat different copies of each gene within the genome of an individual as different alleles of the gene obtained from different individuals. Mathematically, this is not a safe practice, but here we set out to explore the performance of the method when the methods are used in such a way.

Figure 5 shows the frequency of correct inferences of the phylogenetic network topologies by these two methods on data generated under Scenario 1 of figure 4.

**Figure 5:**
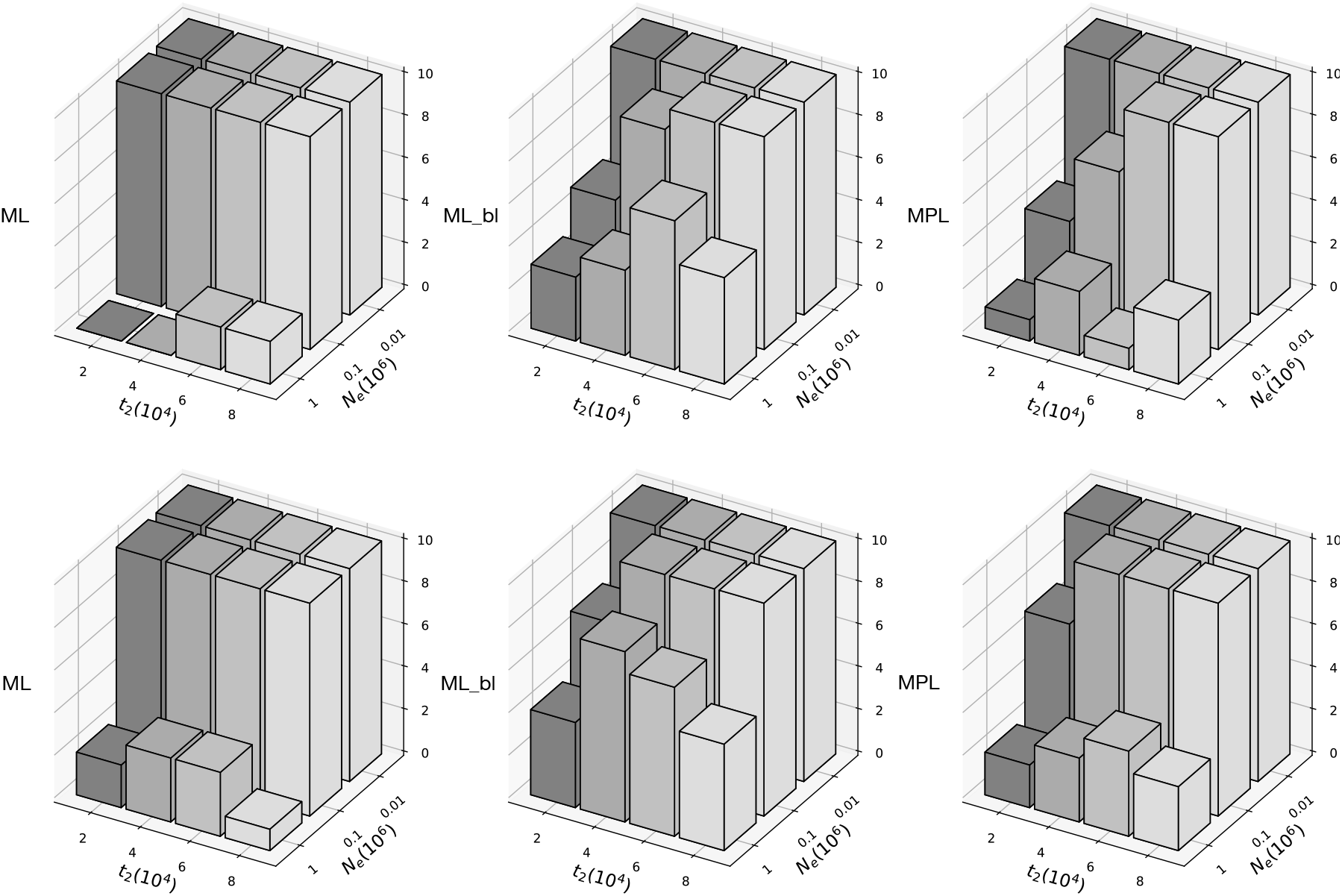
The number of correct estimates of networks with specified hybridization on simulated data: MSNC-based methods. Each bar represents the number of correctly inferred networks out of 10. The true phylogenetic network is the one in Scenario 1. Top row: Each data set consists of 50 gene trees. Bottom row: Each data set consists of 100 gene trees.

There are several points to make here. ML on gene tree topologies alone is the most accurate of all three methods. In particular, ML inferred the correct phylogenetic networks under all values of *t*_2_ except for the largest population size setting. And this was achieved when using both 50 and 100 gene trees in the input. The population size controls the amount of ILS in the data. For the largest value of *N_e_*, the amount of ILS is largest, resulting in coalescence patterns of A, B, and T lineages that mislead the inference. Increasing the number of gene trees from 50 to 100 helped improve the results, but not significantly. When using gene trees with branch lengths as input, ML has perfect accuracy for the smallest population size setting (very little ILS). However, for a population size that is one order of magnitude larger, the performance now is not as good as when using gene tree topologies alone. While this might seem unintuitive (as gene trees with branch lengths provide more information to the inference method), an increased level of ILS forces the inference method to change the topology in order to satisfy the coalescent time constraints presented by the input gene trees. However, this decrease in accuracy is very slow that even when the population size is set to the largest possible value in these simulations, the performance is much better than that of ML on gene tree topologies alone.

MPL had the lowest accuracy in this category of methods. This makes sense as pseudo-likelihood is an approximation of the full likelihood designed to speed up the calculations. Indeed, for larger data sets, running ML, especially when using gene tree topologies alone, is infeasible, whereas MPL could be used.

We observe very similar trends on data from Scenarios 2 and 3 (supplementary figs. S1–S3, Supplementary Material online), though the accuracy on Scenario 2 was highest. The reason for this is that in Scenario 2, even when *t*_2_ is small, the parents of the hybrid are much more diverged than in the other two scenarios, which involve two sister taxa.

As counting only correctly inferred networks is a strict measure of accuracy, we also quantified the true positive and negative rates of all inferences (both values are 1 when the network is correctly inferred). The results are shown in figure 6 for all three scenarios combined. These results show clearly that, except for the largest value of *N*_e_, ML, ML_bl, and MPL have very good accuracy of the inferences, even when not inferring the true network. For the largest population size, the accuracy of all three methods is over 85% when 100 gene trees are used. In other words, while not many networks were inferred perfectly for the largest population size, the accuracy of the inferred networks is very high.

**Figure 6:**
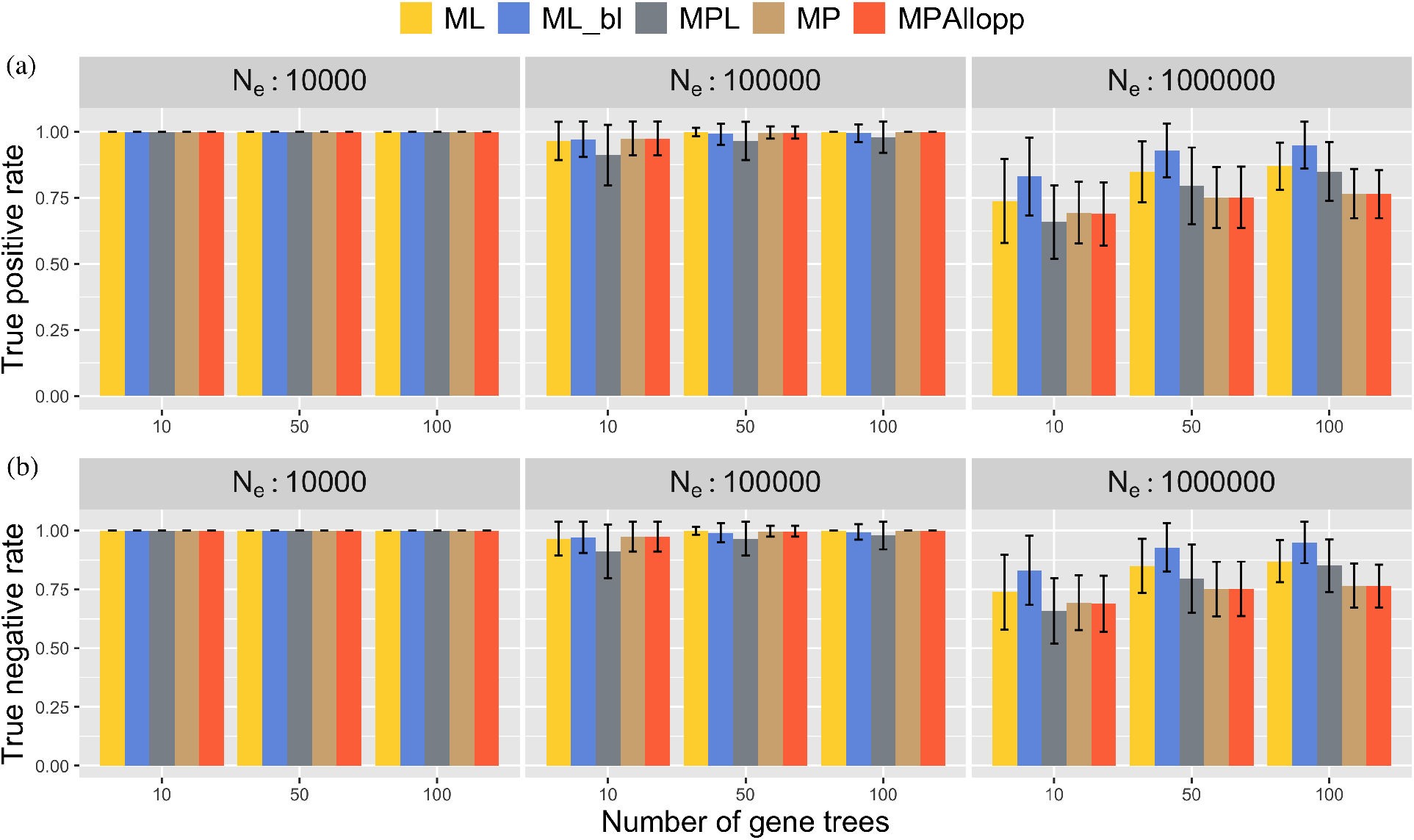
Accuracy of inference results on simulated data. True positive rates (a) and true negative rates (b) are shown for 10 replicate data sets and all three scenarios and all values of *t*_2_ combined.

It is important to note that while inference under the MSNC could result in the correct phylogenetic network topology, several of the branch lengths could be inferred incorrectly. Consider the scenario of figure 1c. Since inference under the MSNC assumes that *x*_1_ and *x*_2_ are two alleles of species *X*, and the same for the pair (*y*_1_,*y*_2_) and the pair (*z*_1_, *z*_2_), the lengths of all four branches in the subtree ((*X, Y*), *Z*) would be underestimated to account for the absence of coalescence events on these branches. Indeed, we inspected the length of the branch connected to T in the ML inferred networks and found out that 178 out of 180 inferred networks have shorter branch length than the ground truth. Moreover, all the ML correctly inferred networks have shorter branch lengths, which is consistent with our expectation.

In other words, while both ML and MPL happen to provide good results in these simulations in terms of the phylogenetic network topology, they did so at the expense of the branch lengths. For example, for Scenario 1, the coalescent times of the homoeologous alleles from the tetraploid T have to be more ancient than the divergence time between species A and B. That is to say, no homoeologs could coalesce along the two horizontal edges or the branch connected to T. So, even when the tetraploid T was correctly inferred as the hybrid species, ML worked by forcing the age of the hybridization to be zero.

#### 3.2 Performance of Parsimony Methods

Next, we ran `InferNetwork_MP`, which implements the maximum parsimony method of [35] (labeled MP below), and `InferNetwork_MP_Allopp`, which is the new method described above (labeled MPAllopp below). Results in terms of the frequency of correctly inferred networks for Scenario 1 are shown in figure 7.

**Figure 7:**
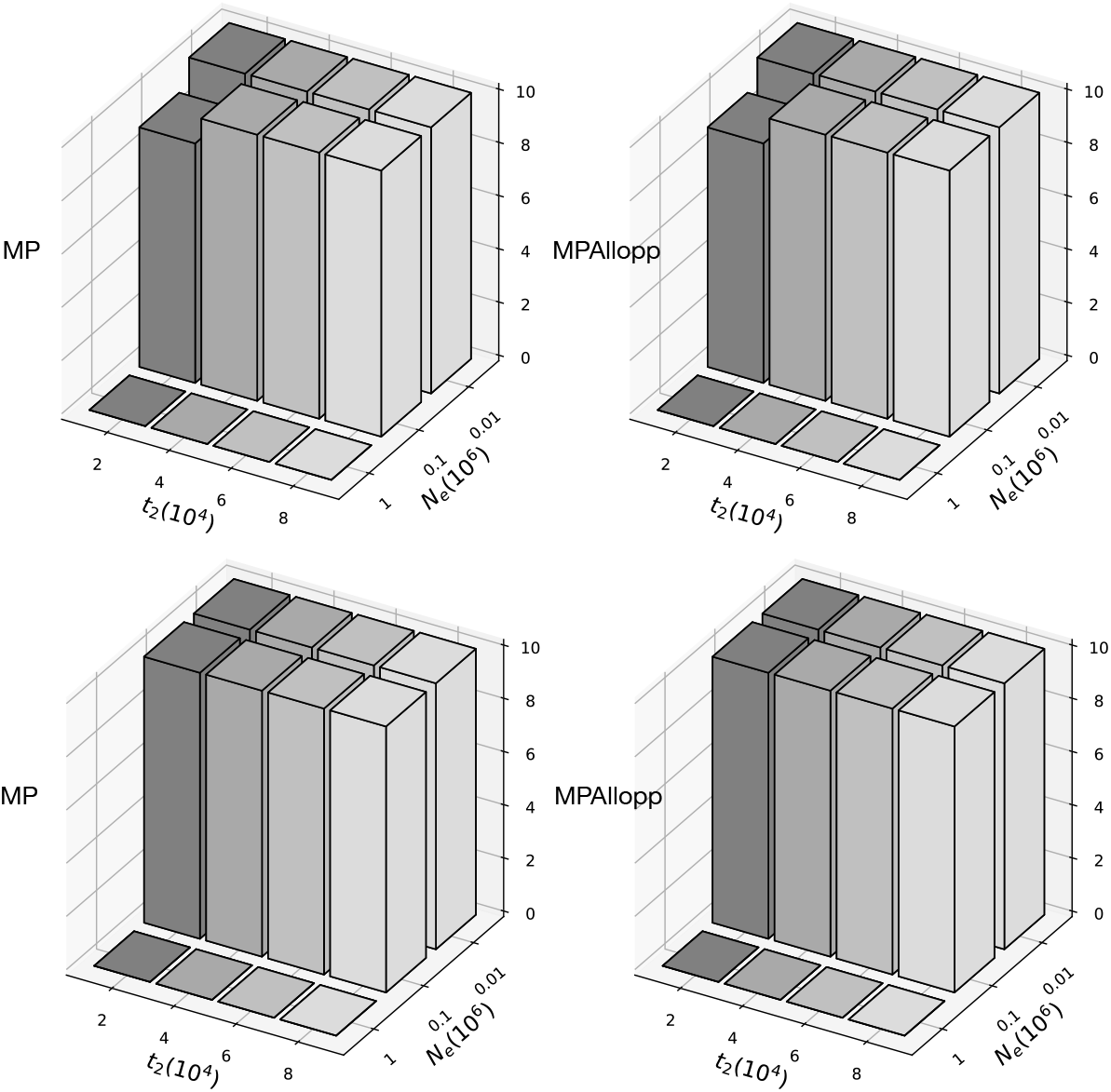
The number of correct estimates of networks with specified hybridization on simulated data: Parsimony methods. Each bar represents the number of correctly inferred networks out of 10. The true phylogenetic network is the one in Scenario 1. Top row: Each data set consists of 50 gene trees. Bottom row: Each data set consists of 100 gene trees.

As the figure shows, both methods have almost identical performance, and both are very similar to ML above. Furthermore, as with the methods above, we observe very similar trends on data from Scenarios 2 and 3 (supplementary figs. S4 and S5, Supplementary Material online), though the accuracy on Scenario 2 was highest. Figure 6 shows that in terms of accuracy of the inferred networks the two methods are identical.

However, as with ML and MPL, while MP inferred the network topology with the same accuracy as the new method, this does not mean that MP is equally appropriate as MPAllopp. We illustrate this through an example. Consider the phylogenetic network of figure 8a. The model network and parameters were obtained from [11], where the population size was 10^5^ individuals, and the mutation rate was 8 × 10^−8^ per site per generation. 30 gene trees were simulated (supplementary fig. S6, Supplementary Material online), using the simulation program AlloppDT of [11]. Then `InferNetwork_MP` and `InferNetwork_MP_Allopp` were used to infer the species network with one reticulation node. As shown in figure 8, MP failed to infer an evolutionary history with hybridization, returning a species tree instead. MPAllopp, on the other hand, made correct inference with a MPAllopp score of 108. To ensure that the failure of MP on this data set was not due to failed search, we calculated the MP score of the true network topology using the command `DeepCoalCount_network`, and we found that the true topology has a score of 280, which is worse than that of the inferred tree, which is 245.

**Figure 8:**
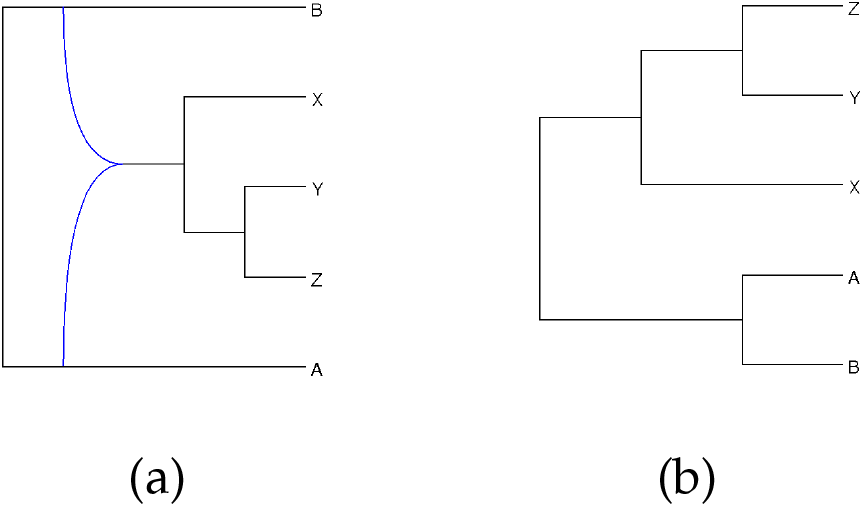
An example scenario where MP fails to recover allopolyploidization when it is present. (a) The true evolutionary history. Here, a hybridization occurred between these two diploid species A and B, forming an allotetraploid from which the XYZ clade descended. (b) The phylogenetic network inferred by MP.

#### 3.3 Distinguishing Auto- and Allo-polyploidy Under the Parsimony Criterion

We illustrate here how the parsimony criterion can be used to determine whether auto- or allo-polyploidization took place. As we discussed above, in the former case, the evolutionary history of the species should be a tree, not a network, whereas in the latter case it should be a network. Therefore, when running MP or MPAllopp on data including autopolyploids, the method would find that a tree fits the data equally well to a network (barring weak signal) and return a tree. In the presence of allopolyploids, the method would find that a network fits the data better than a tree.

To demonstrate the utility of MP for identifying autopolyploidy, we applied MP to an example data set obtained from https://gwct.github.io/grampa/example1.html. This data set includes 1000 gene trees whose underlying species phylogeny is shown in figure 9. We ran MP, setting the number of allowable reticulations to 1, and MP accurately recovered the underlying singly-labeled species tree even though hybridization was allowed, as shown in figure 9.

**Figure 9:**
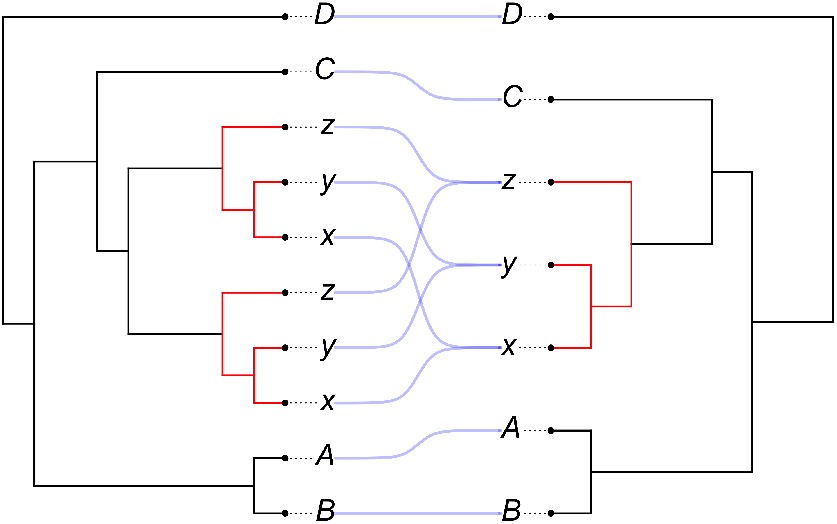
An example data set involving autopolyploidy. Here, a WGD event took place at the MRCA of ((x,y),z). Left: the species MUL-tree used to simulate the gene trees in the presence of autopolyploidy. Right: the species tree topology inferred by MP, which is identical to the true singly-labeled topology.

#### 3.4 Running Times of the Methods

The average running times across all conditions and data sets in CPU minutes of ML, ML_bl, MPL, MP and MPAllopp are 1503.60, 23.87, 256.91, 19.93, and 19.95, respectively. Figure 10 gives a more detailed view of the running times as a function of the two parameters that affect running time most, namely the number of gene trees in the input and the effective population size.

**Figure 10:**
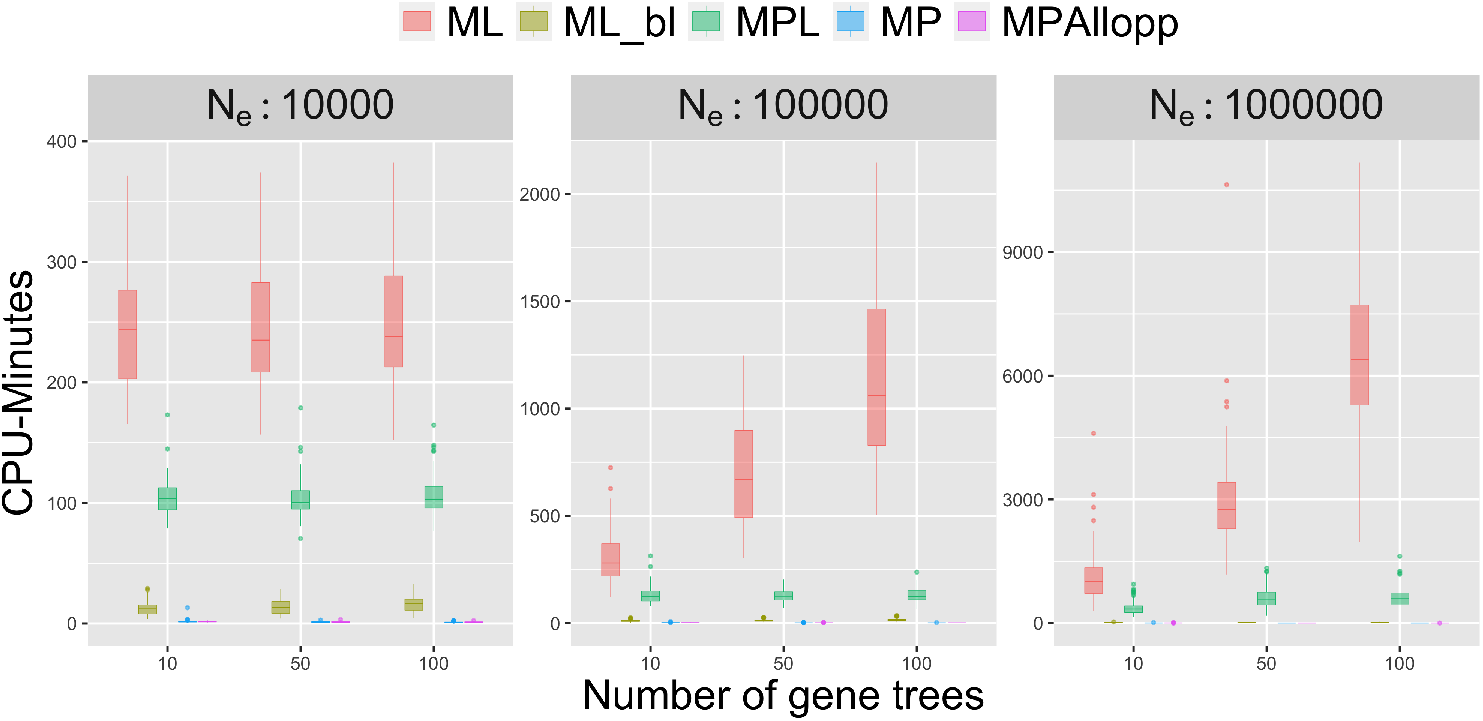
Running times of the five methods in CPU minutes. Results are shown for 10 replicate data sets. Note: The scales of the y axes in the three panels differ significantly.

As the population size increases, the heterogeneity between the gene trees and the phylogenetic network, as well as among the gene trees themselves, increases due to ILS. This in turn results in increased running times of computing the gene tree probabilities. This explains the differences in running times of ML across the three population sizes. Furthermore, this also explains the sharper increase in ML’s running time as the number of gene trees increases in the case of the larger population sizes than in the smaller ones.

When the branch lengths of the gene trees are used (ML_bl), the likelihood calculations are cut significantly because the number of possible coalescent histories decreases sharply [37].

As maximum pseudo-likelihood relies on frequencies of triplets of taxa [38], the running times are hardly impacted by the population sizes and are impacted linearly in terms of the number of gene trees.

The running times of both parsimony methods are negligible as these methods do not estimate branch lengths of the networks.

#### 3.5 Analysis of a Biological Data Set

We ran `InferNetwork_MP_Allopp` on the biological data set in [22], which consists of diploid, tetraploid and hexaploid representatives of the genus *Leucanthemopsis* (Compositae, Anthemideae). The data set consists of four nuclear single-copy genes and two plastidic intergenic spacer regions. We downloaded the sequences from the GenBank database using the accession numbers provided in [22], and aligned the sequences using MAFFT version 7.453 [13]. For each locus, we ran BEAST version 1.10 [26] to infer a sample of trees, based on which we computed the maximum clade credibility tree, resulting in five inferred gene trees, as one tree was inferred on the two intergenic spacer regions (supplementary figs. S7–S11, Supplementary Material online).

In our analysis, we set the maximum number of allowed reticulations to 0, 1, 2 and 3, and with 0, 1 (the hexaploid *L. alpina* subsp. *cuneata*) or 2 (the hexaploid *L. alpina* subsp. *cuneata* and the hexaploid *L. longipectinata*) specified hybrid species.

We obtained the optimal results of MPAllopp after 20 runs of search, as are shown in Table 1. As the table shows, when the hybrid species were specified, an optimal network was obtained even when allowing extra hybridizations to be added. Based on these results, the optimal network with a single hybridization is the one with the hexaploid *L. alpina* subsp. *cuneata* specified as hybrid. The optimal network with two hybridizations is the one with the hexaploid *L. alpina* subsp. *cuneata* and the hexaploid *L. longipectinata* specified as hybrids.

**Table 1:** Results of InferNetwork_MP_Allopp on the *Leucanthemopsis* data.

| InferNetwork_MP_Allopp |  |  |
| --- | --- | --- |
| #Specified hybrid species | Maximum #reticulations | MPAllopp score |
| 0 | 0 | 241 |
| 1 | 1 | 233 |
| 1 | 2 | 233 |
| 1 | 3 | 233 |
| 2 | 2 | **227** |
| 2 | 3 | **227** |

The optimal network with two hybrids is shown in figure 11. This network shows that that an allotetraploid was created through allopolyploidzation, and this allotetraploid later hybridized with a diploid species (*Castrilanthemum debeauuxu*), giving rise to a hexaploid that is the ancestor of the two hexaploid species *L. alpina* subsp. *cuneata* (6x) and *L. longipectinata* (6x).

**Figure 11:**
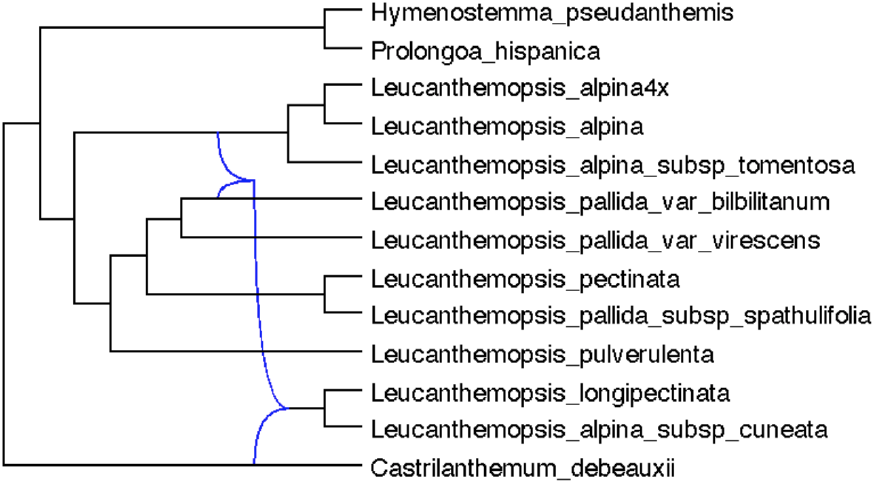
Inference results using the MPAllopp criterion on the example data set of *Leucan-themopsis* representatives in [22]. The optimal network with a MPAllopp score of 227 when the maximum number of reticulations was set to 2 and *L. alpina* subsp. *cuneata* (6x) and *L. longipectinata* (6x) were specified to be the hybrid species.

The tetraploid *L. alpina* was found to be an autopolyploid. Even though the *L. alpina* subsp. *cuneata* (6x) is one of the intraspecific taxa of the widespread *L. alpina* Heywood, it is postulated to have a closer relationship with other Iberian species. This might be due to the spatial locality, as the *L. alpina* subsp. *cuneata* (6x) lives in Northern Spain.

Finally, the network we inferred and reported in figure 11 above is different from the network inferred and reported by [22]. Using the DeepCoalCount_AlloppNet we calculated the MPAllopp score of the network of [22] using the gene trees we inferred. The score is 269, which is much worse than the tree we inferred (which has a score of 241). It is important to note here that [22] used different gene tree estimates. When we used the gene tree estimates reported in the original study, we found that the MPAllopp score of the network of [22] was 154, whereas the network we reported in figure 11 had a score of 134. To summarize, regardless of the gene trees used (which differ in the way they were inferred), the network we report here is a better network in terms of the MPAllopp score.

### 4 Concluding Remarks

In this paper, we introduced a new maximum parsimony method for inferring phylogenetic networks from gene tree topologies while accounting for polyploidy and incomplete lineage sorting simultaneously. The method employs a heuristic search for walking the network space while evaluating the parsimony score on the MUL-tree representation of the network.

A question that begs to be answered is: If MUL-trees can be treated as equivalent to phylogenetic networks, then what is the gain from using the latter model? The lack of a one-to-one mapping between MUL-trees and phylogenetic networks notwithstanding [9, 43], “seeing” the polypoloid hybridization events in a MUL-tree is possible only for simplistic scenarios: a small number of taxa, a small number of hybridization events, a small number of gene extinctions, and, most importantly, the absence of confounding factors such as ILS (figure 1). Indeed, the parsimony algorithms and methods of [8, 29] do not account for ILS. Identifying the hybridization events computationally is the task of turning the MUL-tree into a phylogenetic network after a MUL-tree is inferred from the gene trees. Therefore, our method searches the phylogenetic network space directly by applying sub-network transfer operations on networks, so that the inference result is a network, rather than a MUL-tree.

As discussed in [2], statistical modeling of phylogenetic networks with polyploid hybridization is very complex. We believe that devising stochastic models and inference methods for restricted classes of polyploids, as in [11, 24], is most likely the way to make progress in this area.

In the last several years, there has been work on combining the multispecies coalescent model with a birth-death model of gene duplication and loss [34, 25, 5, 14]. Most recently, [6] introduced a model that unifies the multispecies network coalescent and a birthdeath model thus allowing for simultaneous modeling of incomplete lineage sorting, gene duplication and loss, and diploid hybridization. These works could be relevant for further advances in modeling polyploid hybridization in phylogenomic inference.

Statistical inference of phylogenetic networks is computationally much more demanding than inference of trees, severely limiting the sizes of data sets that can be analyzed with phylogenetic network methods. One approach to handling larger data sets is to analyze smaller subsets of the data (subsets in terms of taxa). This approach could be automated, as in [40] for example, but this requires developing methods for accurately estimating small networks with their evolutionary parameters and for merging these small networks into a network on the full data set. We view this as an essential direction for future research for phylogenetic network inference on large data sets involving polyploids to become feasible.

## Acknowledgments

This work was supported by grants CCF-1514177, CCF-1800723, and DMS-1547433 from the National Science Foundation provided to L.N.

## Notes

### Competing Interest Statement

The authors have declared no competing interest.

