## Supplementary Material for "Maximum Parsimony Inference of Phylogenetic Networks in the Presence of Polyploid Complexes"

Zhi Yan, Zhen Cao, Yushu Liu, and Luay Nakhleh
*Department of Computer Science*
*Rice University*

- Accuracy of inferred species networks on the 5-taxon simulated data sets from true gene tree topologies using ML (on gene tree topologies and on gene trees with branch lenghts), MPL, MP, and MPAllopp are shown in Figs. S1—S5. In all these analyses, the polyploid was specified to the method.
- The gene trees simulated by AlloppDT are shown in Fig. S6.
- Estimated gene trees for the *Leucanthemopsis* data are shown in Figs. S7—S11.

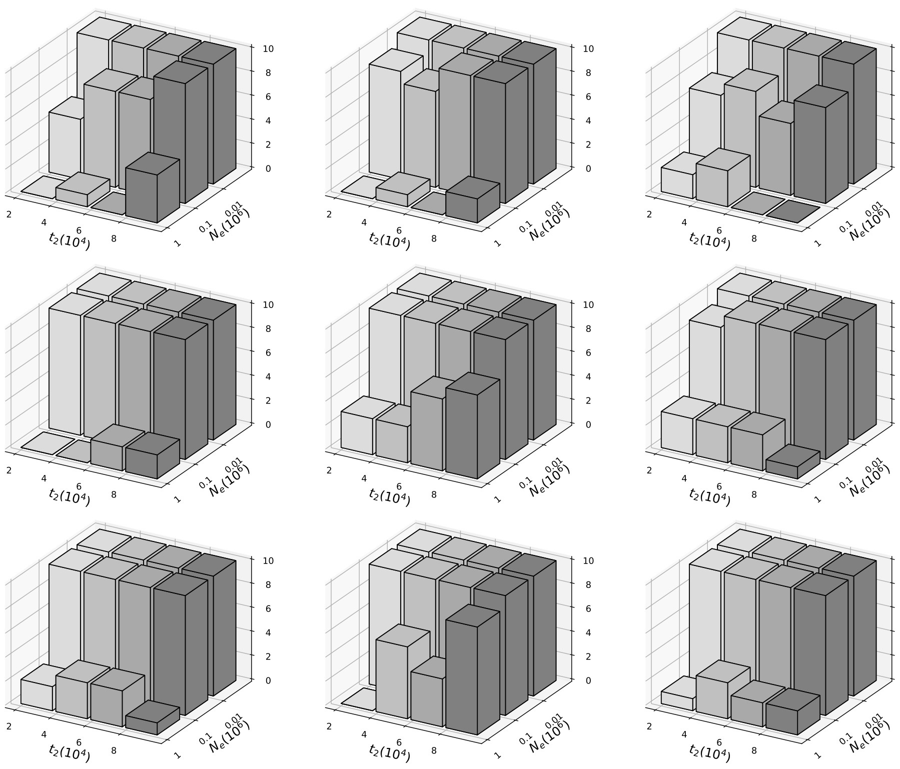


Figure S1: **The number of correct estimates of networks under the ML criterion when using gene tree topologies.** From left to right, columns represent scenario 1, scenario 2 and scenario 3, respectively. From top to bottom, rows represent 10, 50, 100 gene trees respectively, each of which has 10 replicates. In each subfigure, the x-axis shows branch length $t_2$, varying between 20000, 40000, 60000 and 80000; Y-axis shows the population size $N_e$, including 10000, 100000, 1000000; Z-axis shows the number of correct estimates, varying from 0 to 10, where 0 means all 10 replicates of gene trees inferred wrong networks, while 10 means all 10 replicates inferred correct networks.

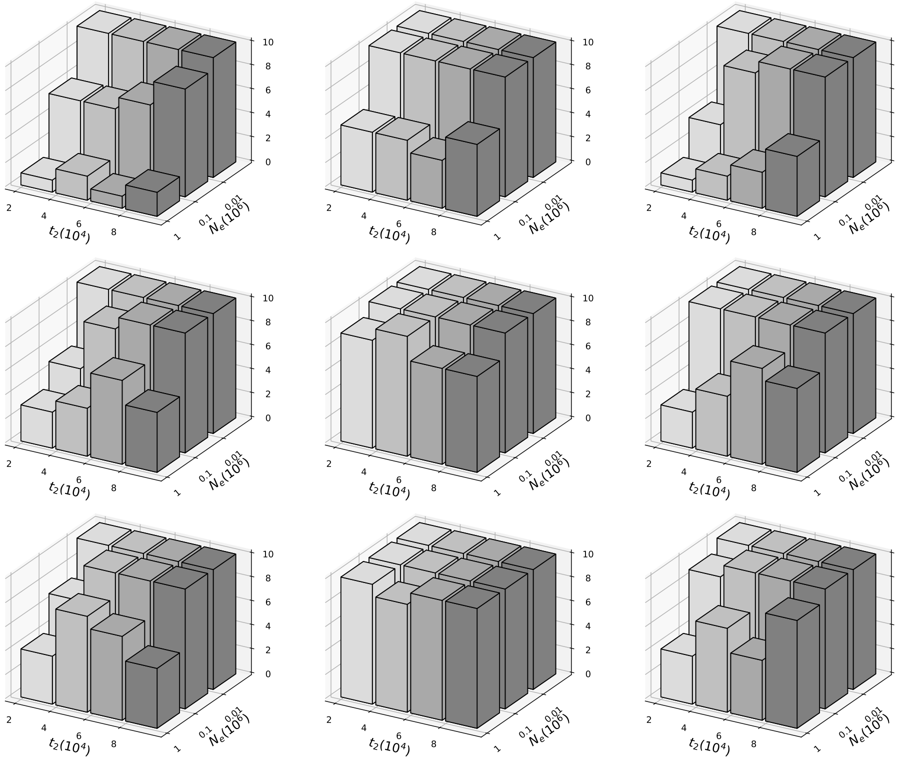

Figure S2: **The number of correct estimates of networks under the ML criterion when using gene trees with branch lengths.** From left to right, columns represent scenario 1, scenario 2 and scenario 3, respectively. From top to bottom, rows represent 10, 50, 100 gene trees respectively, each of which has 10 replicates. In each subfigure, the x-axis shows branch length $t_2$, varying between 20000, 40000, 60000 and 80000; Y-axis shows the population size $N_e$, including 10000, 100000, 1000000; Z-axis shows the number of correct estimates, varying from 0 to 10, where 0 means all 10 replicates of gene trees inferred wrong networks, while 10 means all 10 replicates inferred correct networks.

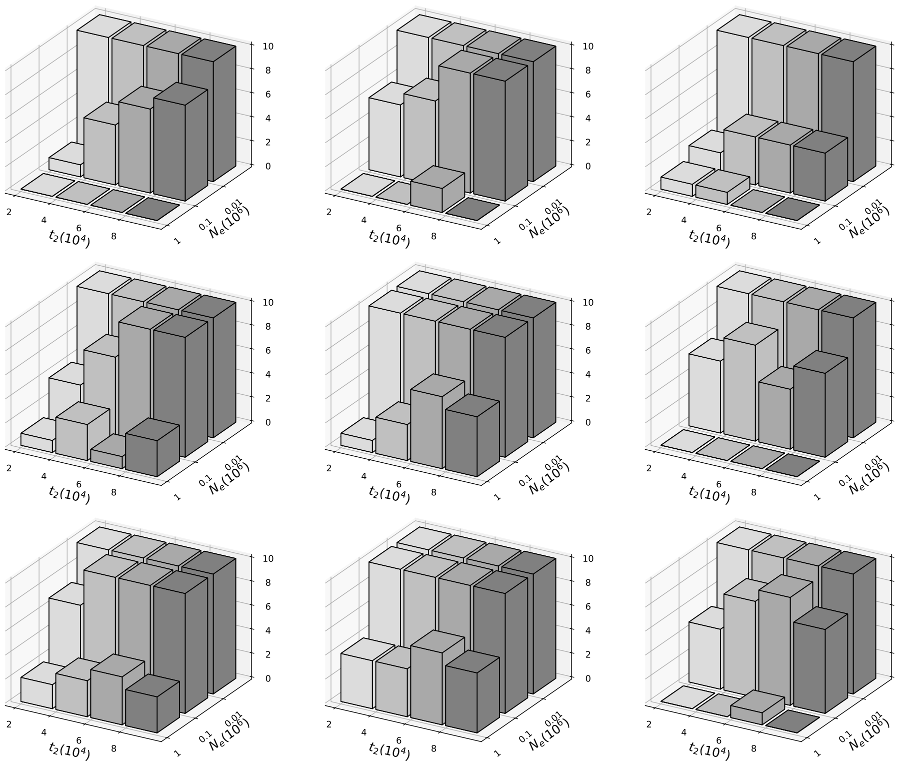


Figure S3: **The number of correct estimates of networks under the MPL criterion.** From left to right, columns represent scenario 1, scenario 2 and scenario 3, respectively. From top to bottom, rows represent 10, 50, 100 gene trees respectively, each of which has 10 replicates. In each subfigure, the x-axis shows branch length $t_2$, varying between 20000, 40000, 60000 and 80000; Y-axis shows the population size $N_e$, including 10000, 100000, 1000000; Z-axis shows the number of correct estimates, varying from 0 to 10, where 0 means all 10 replicates of gene trees inferred wrong networks, while 10 means all 10 replicates inferred correct networks.

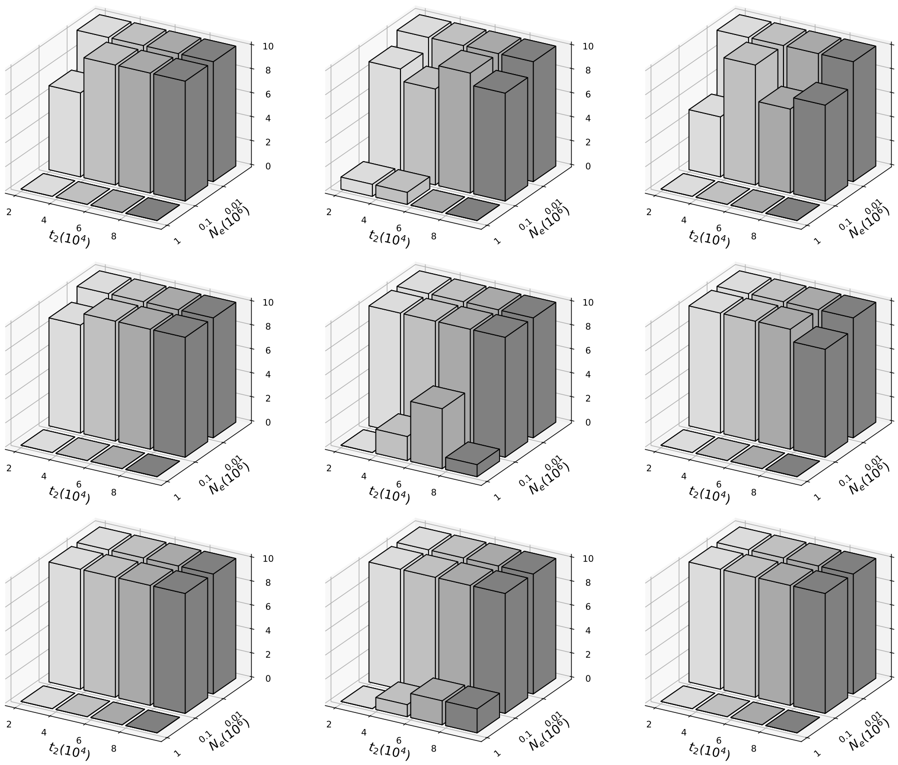


Figure S4: **The number of correct estimates of networks under the MDC criterion.** From left to right, columns represent scenario 1, scenario 2 and scenario 3, respectively. From top to bottom, rows represent 10, 50, 100 gene trees respectively, each of which has 10 replicates. In each subfigure, the x-axis shows branch length $t_2$, varying between 20000, 40000, 60000 and 80000; Y-axis shows the population size $N_e$, including 10000, 100000, 1000000; Z-axis shows the number of correct estimates, varying from 0 to 10, where 0 means all 10 replicates of gene trees inferred wrong networks, while 10 means all 10 replicates inferred correct networks.

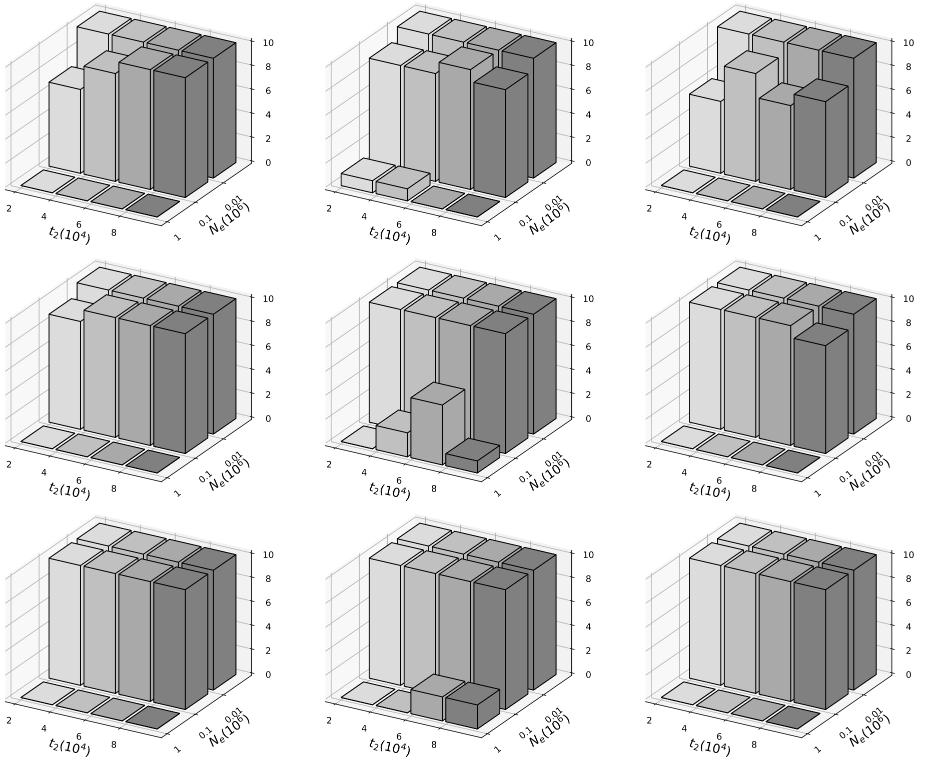


Figure S5: **The number of correct estimates of networks under the MPAllopp criterion.** From top to bottom, rows represent 10, 50, 100 gene trees respectively, each of which has 10 replicates. In each subfigure, the x-axis shows branch length $t_2$, varying between 20000, 40000, 60000 and 80000; Y-axis shows the population size $N_e$, including 10000, 100000, 1000000; Z-axis shows the number of correct estimates, varying from 0 to 10, where 0 means all 10 replicates of gene trees inferred wrong networks, while 10 means all 10 replicates inferred correct networks.

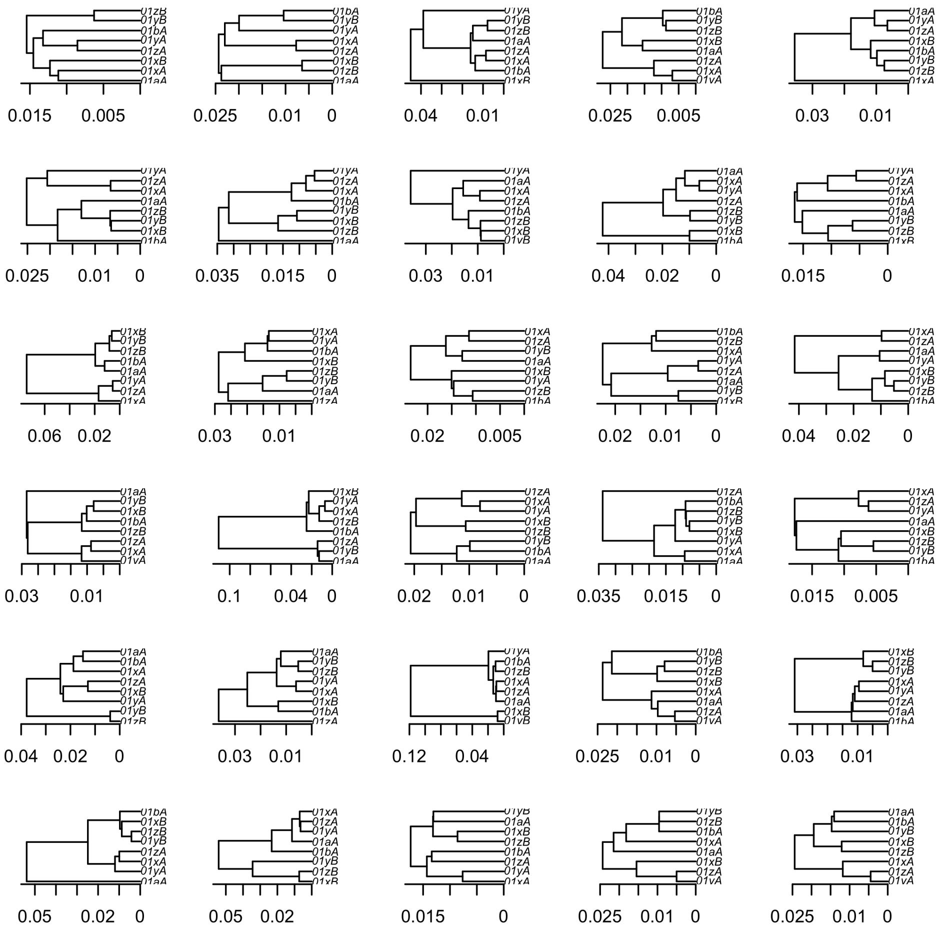

Figure S6: **Simulated gene trees by AlloppDT**. Branch lengths are in units of expected number of substitutions.

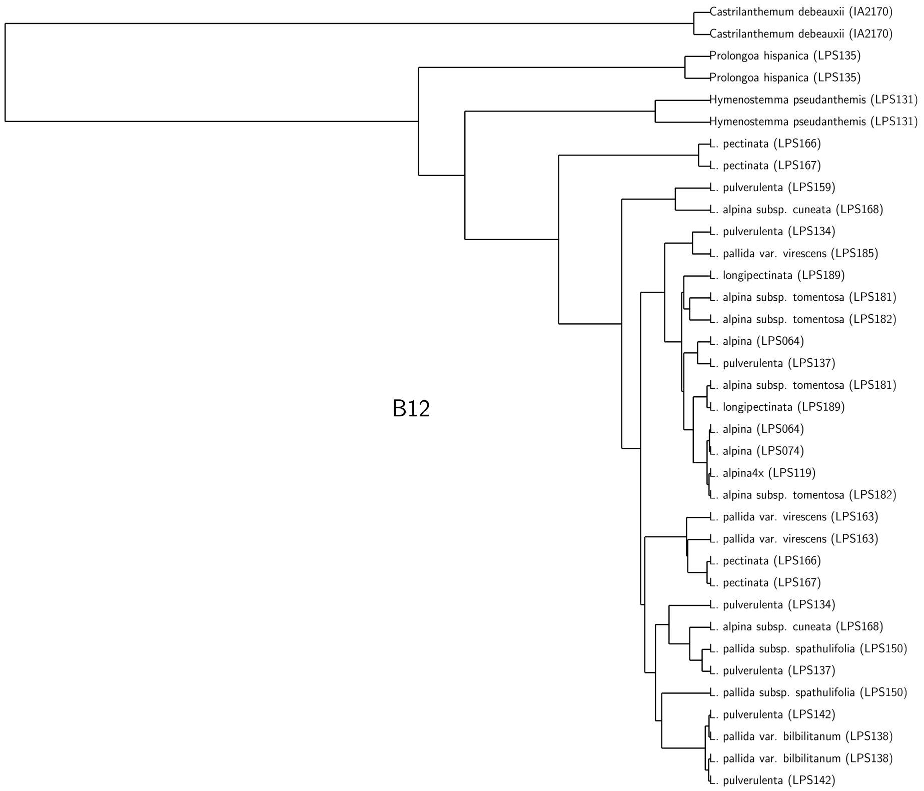


Figure S7: **Maximum clade credibility gene tree based on sequence variation of marker B12**.

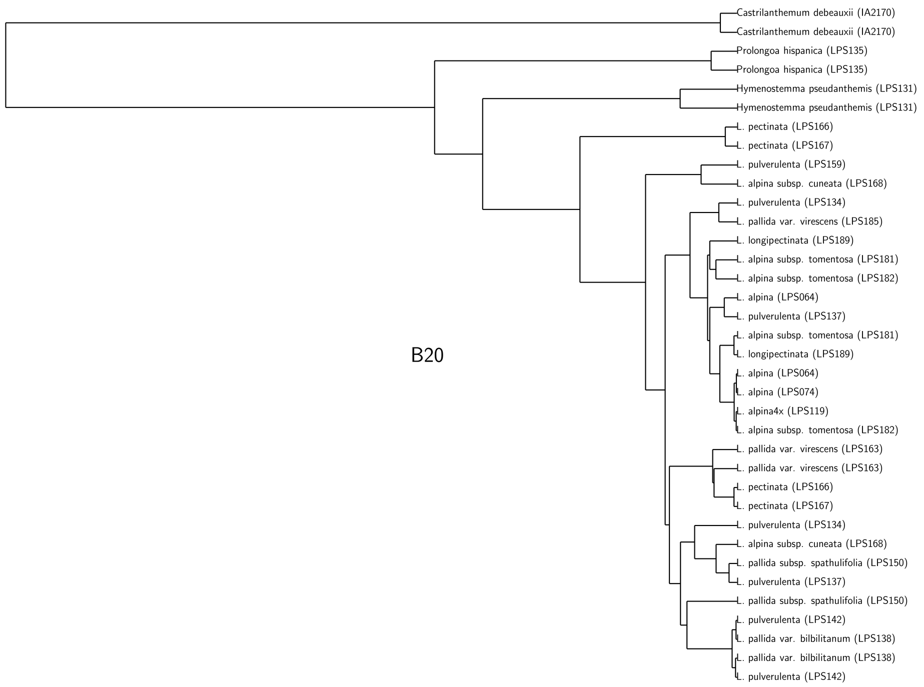


Figure S8: **Maximum clade credibility gene tree based on sequence variation of marker B20**.

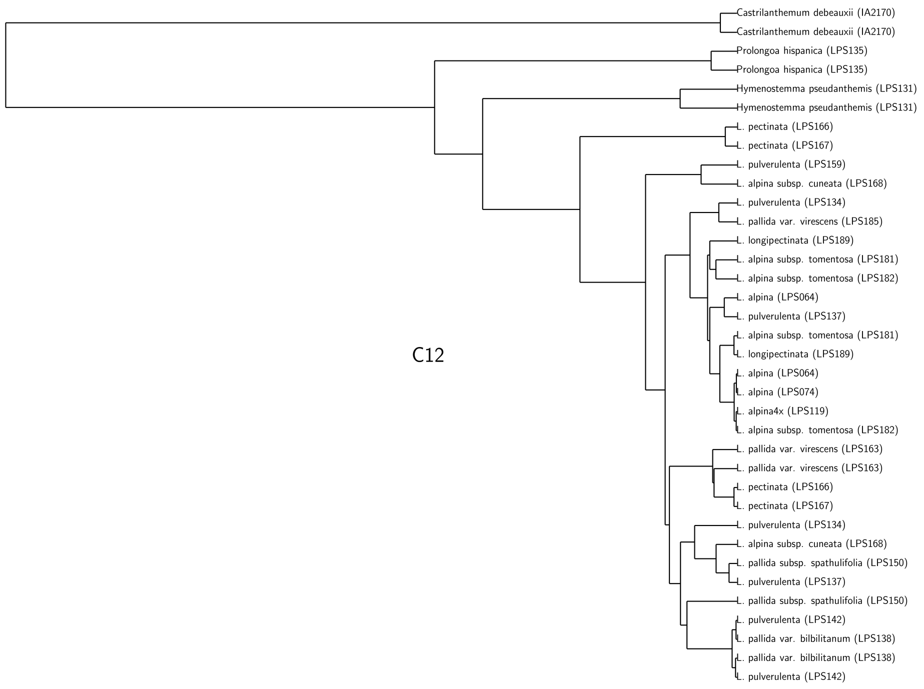


Figure S9: **Maximum clade credibility gene tree based on sequence variation of marker C12**.

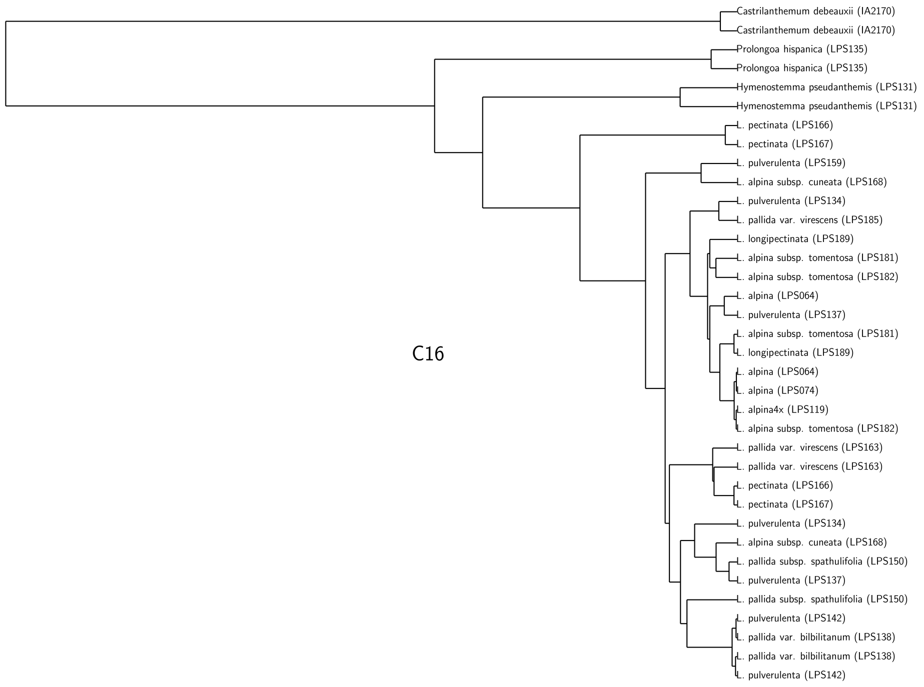


Figure S10: **Maximum clade credibility gene tree based on sequence variation of marker C16**.

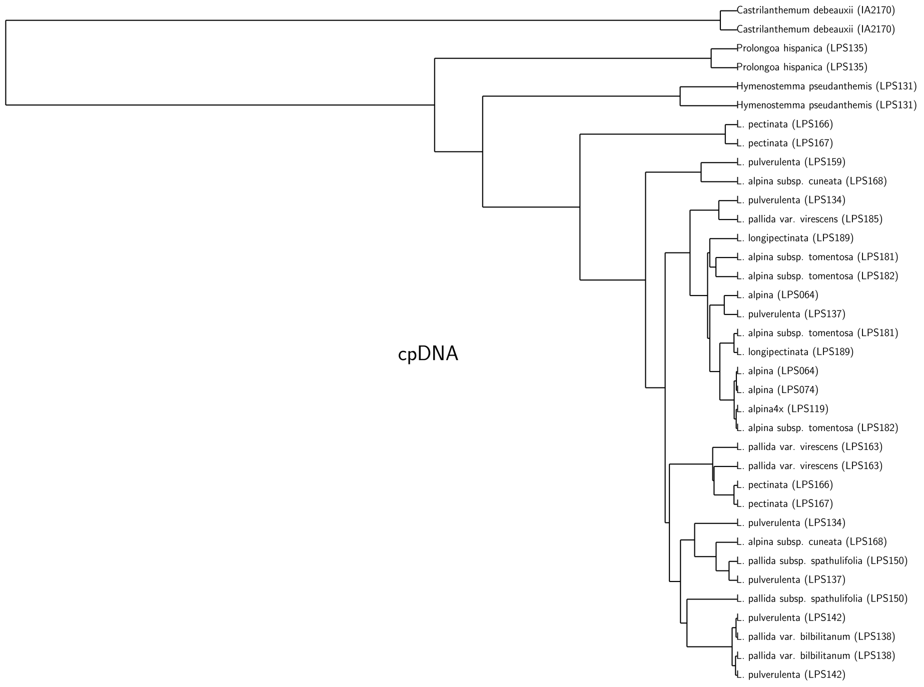


Figure S11: **Maximum clade credibility gene tree based on sequence variation of the two intergenic sparcer regions of the chloroplast genome (*psb-A-trnH* and *trnC-petN*)** .
